# Alpha and theta oscillations contribute to attribute regulation in dietary decision making under self-control

**DOI:** 10.1101/2020.07.09.195958

**Authors:** Azadeh HajiHosseini, Cendri A. Hutcherson

## Abstract

How do different cognitive self-regulation strategies alter attribute value construction (AVC) and evidence accumulation (EA)? We recorded EEG during food choices while participants responded naturally or regulated their choices by focusing on healthy eating or decreasing their desire for all food. Using a drift diffusion model (DDM), we predicted the time course of neural signals associated with AVC and EA. Results suggested that suppression of frontal and occipital alpha power matched model-predicted EA signals: it tracked the goal-relevance of tastiness and healthiness attributes, predicted individual differences in successful down-regulation of tastiness, and conformed to the DDM-predicted time course of EA. We also found an earlier rise in frontal and occipital theta power that represented food tastiness more strongly during regulation, and predicted a weaker influence of food tastiness on behaviour. Our findings suggest that different regulatory strategies may commonly recruit theta-mediated control processes to modulate the attribute influence on EA.

## Introduction

The modern world represents many challenges for the health-conscious consumer. Tasty but unhealthy options abound at grocery stores, restaurants, and social events, with a ubiquity that often confounds even the most disciplined among us. In the face of temptation, people frequently attempt to regulate their responses to food, oftentimes using different strategies. For example, in a survey of obese participants trying to lose weight (Nicklas, Huskey, Davis, & Wee, 2012), 65% reported trying to eat less food overall, and over 40% reported trying to switch the types of foods they ate towards less calorie-dense options. Yet such strategies are at best only partially successful at helping people lose weight (Nicklas et al., 2012). In the face of these results, a major question concerns not only *how* people regulate their choices regarding foods, but also why some are successful while others are not. Here, we took a computational decision neuroscience approach to this problem, using recently developed computational models of choice to ask two questions. First, how do the computations contributing to a choice change when people regulate their behaviour in different ways? Second, which of these changes predict success or failure in altering food consumption?

Recent neuroeconomic theories of decision making suggest that value-based decisions can be captured by a process involving several distinct computations, including both attribute valuation and evidence accumulation (Rangel, Camerer, & Montague, 2008). For example, when we decide what to eat for lunch, we might consider attributes like how tasty or healthy a food is, our dieting goals, current hunger levels, and so forth. Each of these attributes constitutes evidence for or against eating the food, with different attributes given differential weight depending on a person’s momentary goals. To choose, current models assume that noisy neural representations of this attribute-based evidence may serve as the input to a process of sequential evidence accumulation (EA) over time until sufficient evidence has accumulated to pass a decision threshold (Forstmann, Ratcliff, & Wagenmakers, 2016; Gold & Shadlen, 2007).

Sequential accumulation models are both behaviourally and biologically plausible (Forstmann et al., 2016) and appear to capture patterns of choice, response time and neural activity with remarkable accuracy. For example, in the perceptual decision making domain, electroencephalogram (EEG) studies have shown that decrements in the power of alpha (9-12 Hz) and theta (4-8 Hz) oscillations correlate with the predicted build-up of evidence (vanVugt, Simen, Nystrom, Holmes, & Cohen, 2012; Werkle-Bergner et al., 2014). Parallel to this, in the value-based decision making domain, EEG and MEG studies have found that model predictions about the temporal dynamics of EA during decisions correlate with both parietal and frontal theta and alpha (Hunt et al., 2012), and with beta (18-20 Hz) and gamma (40-80Hz) oscillations (Polanía, Krajbich, Grueschow, & Ruff, 2014). Given that value-based decision making engages multiple cognitive processes and involves communication across different regions, these diverse results are perhaps not surprising. However, they raise important questions about what precise computations these oscillatory dynamics represent, and whether and how these signals might be altered by regulatory goals.

These questions are vital for understanding the underlying mechanisms of self-regulation in value-based decision making, since it could target multiple distinct phases of the EA process. Although considerable evidence suggests that self-regulation during decision making modifies choices by changing the influence of attributes on behaviour (Hare, Camerer, & Rangel, 2009; Hare, Malmaud, & Rangel, 2011; Hutcherson, Plassmann, Gross, & Rangel, 2012; Sullivan, Hutcherson, Harris, & Rangel, 2015; Tusche & Hutcherson, 2018), it is unclear how such changes are accomplished neurally. For example, observed decreases in the influence of food tastiness on behaviour could be accomplished either by abolishing attribute representations of tastiness, or by preventing the incorporation of such representations into the EA process.

EEG studies provide some clues in this regard, although they also leave open a number of questions. For example, some research suggests that when instructed to attend to food healthiness, the amplitude of event-related potentials (ERPs) around 300-500 ms post-food-stimulus correlated more strongly with food healthiness. However, it is unclear whether this effect represented changes to early attribute representations or changes to their subsequent weight in the evidence accumulation process. Moreover, while regulation reduced *behavioural* sensitivity to *tastiness* considerations, ERP amplitudes in the same time window correlated with tastiness regardless of regulatory effort (Harris, Hare, & Rangel, 2013). This leaves open the question of how the influence of *tastiness* on behaviour was suppressed.

Given previous research linking oscillatory dynamics to EA (Hunt et al., 2012; Polanía et al., 2014; vanVugt et al., 2012; Werkle-Bergner et al., 2014), we sought to determine whether part of the answer to these questions might lie in the functional role of oscillations in incorporating attribute values into the EA process. More specifically, we used a computational model of choice to first fit the behavioural data, and then simulated the predicted dynamics of both attribute representation and EA processes during food choice, under conditions of natural choice and two distinct regulatory strategies. These results suggested that the two computations might follow distinct temporal trajectories. Then, we sought oscillatory correlates of tastiness and healthiness during self-regulation and asked whether and how the time course of these correlates resembled the simulated computational dynamics of both attribute representation and EA. Finally, we asked whether and how changes in these oscillatory dynamics correlated with individual differences in regulatory success, as measured by changes in model parameters.

## Methods

### Subjects

We recruited 66 participants from the University of Toronto Scarborough research participation system and flyers posted on campus. Five subjects did not complete the task either by choice or due to technical issues. Eleven others were excluded based on our pre-registered exclusion criteria (more than 3 noisy channels in their EEG recording and/or if they made the same choice in more than 90% of trials in our *NATURAL* condition as described under *Task* (Open Science Framework; HajiHosseini & Hutcherson, 2020). Participants were recruited until we obtained our pre-registered target sample size of 50 usable subjects (30 female, age range 17-31). All subjects gave written consent prior to the experiment. The study was approved by the Ethics Board of the University of Toronto. Subjects recruited through the research participation system received course credits while those recruited by flyers received $30 for participation.

### Task

To increase motivation to eat, all participants were instructed to refrain from eating for 3 hours before the start of the experiment. The experiment consisted of three tasks: a *pre-choice rating* task, a *Self-Regulation Choice Task (SRCT)*, and a *post-choice rating* task (Hare et al., 2011; Harris et al., 2013; Hutcherson et al., 2012). Subjects also reported how hungry they were on a scale of 1 to 9 (hunger level) just before the pre-choice rating and after the SRCT. During the task, subjects were seated approximately 75 cm from a 24-inch computer screen.

### Pre-choice rating task

In order to match the values of foods appearing in each condition, we recorded liking ratings for 208 different food stimuli before and after the SRCT. Subjects rated the foods on a scale of 1 to 6 corresponding to *strongly dislike, moderately dislike, slightly dislike, slightly like, moderately like, strongly like*. Stimuli were presented in the center of the screen filling 45% of the screen area. Rating keys were presented under the food picture. Based on these ratings, we selected 180 different foods and divided them into three, roughly equally liked sets of 60 foods each, for use in the SRCT task that followed.

### SRCT

During the SRCT, participants made choices about whether to eat different food stimuli under three different conditions, presented in blocks of 15 trials. 60 unique foods appeared in one of three conditions: focus on healthiness (HEALTH), decrease desire for food (DECREASE), or respond naturally (NATURAL). Subjects completed 36 randomly interleaved blocks (12 blocks of each condition). Each of the 60 food stimuli appeared three times throughout the blocks in a single condition (i.e., 180 trials per condition), in order to avoid confounding memory or value effects across conditions. At the beginning of each block, one of three sets of instructions corresponding to each condition was presented on the screen for 5 seconds:

**NATURAL**: *RESPOND NATURALLY; For the next set of trials, we would like you to RESPOND NATURALLY. Allow any feelings or thoughts you have to come naturally and make whatever choice you most prefer at that moment*.
**DECREASE**: *FOCUS ON DECREASING DESIRE; For the next set of trials, we would like you to DECREASE YOUR DESIRE FOR FOOD. Do whatever you need to in order to decrease your craving for food as you decide what you prefer to do. Then make whatever choice you most prefer at that moment*.
**HEALTH**: *FOCUS ON HEALTHINESS; For the next set of trials, we would like you to FOCUS ON THE HEALTHINESS OF THE FOODS. Consider how healthy the food is as you are deciding what you prefer to do. Then make whatever choice you most prefer at that moment*.

Every trial started with a fixation cross in the center of the screen for 500 ms and was followed by a food stimulus. To minimize eye movements, the food stimulus was centred and filled only 20% of the screen. Subjects pressed one of four response keys (*d*, *f*, *j*, or *k)* on the keyboard to indicate their decision (*Strong No, No, Yes*, or *Strong Yes*) about each food, with right-left order of response counterbalanced across subjects. Participants had 4 seconds to decide. Following response, a randomly jittered 1-2 second inter-trial interval appeared. To ensure that subjects followed the correct instruction throughout every block, we signaled the conditions with a color code: NATURAL = green, DECREASE = yellow, and HEALTH = red. A colored frame appeared around the written instructions at the beginning of each block and the fixation cross at the beginning of each trial appeared in the color corresponding to the condition (Figure 1a).

**Figure 1:**
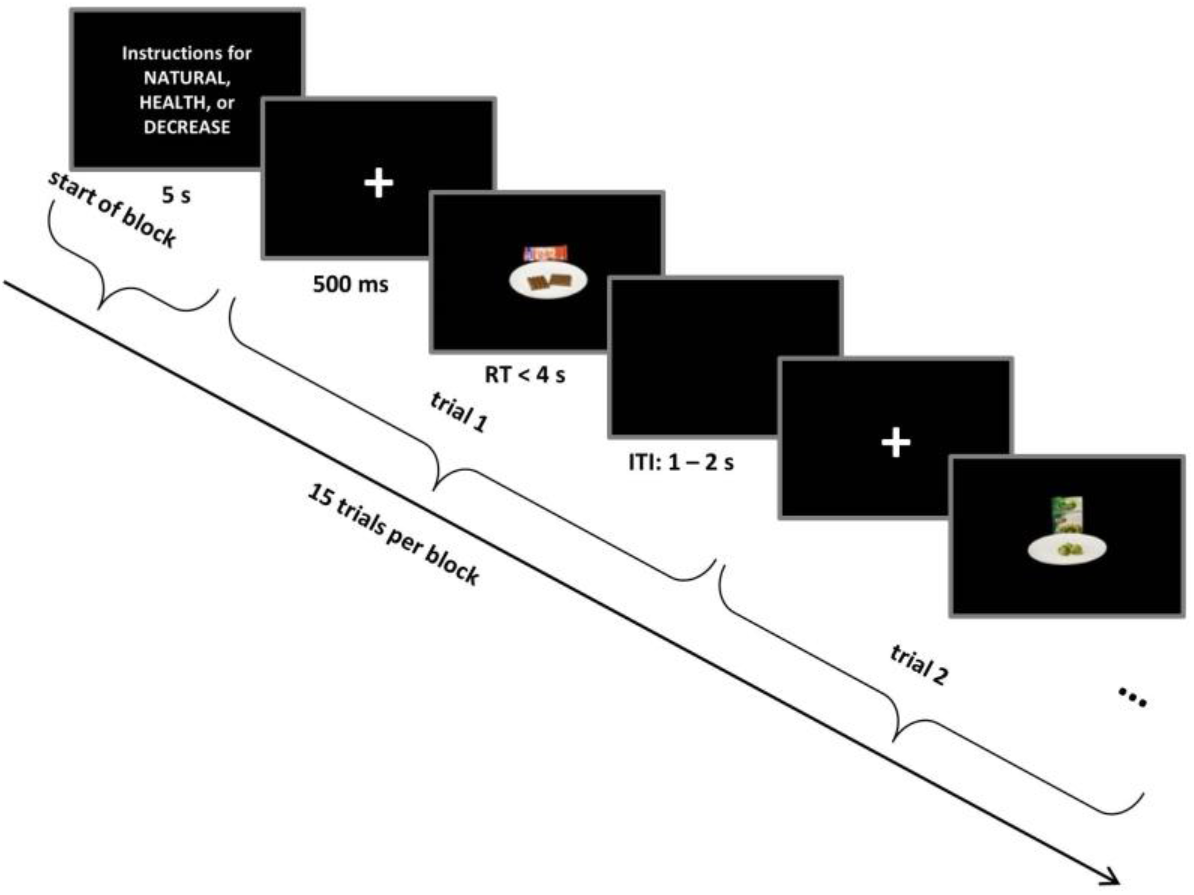
Self-regulation task structure. Trials occurred in interleaved blocks of NATURAL, HEALTH, and DECREASE conditions. Each block started with instructions related to NATURAL or regulation conditions (see Methods) that remained on the screen for 5 seconds followed by 15 choice trials. Every food stimulus was preceded by a fixation cross that was color-coded for the condition.

### Post-choice attribute rating task

In addition to providing a second set of liking ratings for foods after the regulation task, subjects also rated the food stimuli from the SRCT for tastiness and healthiness on a scale of 1 to 6 *(very untasty* to *very tasty* and *very unhealthy* to *very healthy)*. All foods were rated on one attribute, and then on the other, with rating order randomized across subjects. These subjective attribute ratings allowed us to assess how regulatory strategies altered the influence of attribute values on choice.

Following completion of liking, tastiness and healthiness ratings, one trial from the SRCT was randomly selected for implementation and presented on the screen. The subject ate the food if they responded Yes or Strong Yes. Following completion of all experimental tasks, subjects completed a set of questionnaires designed to measure individual differences and real-world dietary behaviour. We wrote the experiment codes in Matlab, using the Psychophysics Toolbox (Brainard, 1997).

### EEG

#### Recording

We used a 64-channel BioSemi ActiveTwo system for recording the EEG during the SRCT, aligned according to the 10-20 system. We also recorded from three external channels, left and right mastoids for offline referencing and an electrode under the right eye to detect eye blinks. Recordings on Biosemi were sampled at 512 Hz, reference free, and later re-referenced to the averaged mastoids upon importing data to EEGLAB (Delorme & Makeig, 2004).

#### Pre-processing

All channel data were high-pass filtered at 0.5 Hz to remove slow drifts, and low-pass filtered at 100Hz to remove high-frequency noise. A notch filter was applied to remove the 60 Hz line frequency. Data were then re-referenced to averaged channels. We identified bad channels by visual inspection and replaced them by interpolation. To centre the data, average amplitude for each channel was subtracted from each data point on that channel. Data were then segmented from 1 sec prior to the presentation of the food stimulus to 1.5 sec after the presentation of the food stimuli. Average amplitude over a baseline window starting from 200ms before the presentation of food stimulus was subtracted from all time points in the segment. Independent Component Analysis (ICA) was then performed on concatenated segments. Components associated with ocular and muscle artefacts or channel noise were identified based on their time and frequency information and removed. Channel data were reconstructed based on the remaining components, and re-segmented into the original segments.

#### Time-frequency analysis

Each trial starting from 1 sec before to 1.5 seconds after the presentation of food stimulus was convolved with a complex Morlet wavelet with a varying number of cycles (ranging from 3 for lower to 10 for higher frequency ranges) to study frequencies from 1 to 80 Hz in 1 Hz steps. The transformed time-series contained a complex number for each time-point, channel, and frequency, *A^jϕt^* where *A* was the power and *ϕ* the phase angle. We normalized the power in each time point (*A_t_*) to average power at a baseline consisting of the 200ms prior to onset of the food stimulus: 10*log_10_((*A_t_* – *A_baseline_*)/ *A_baseline_*). We calculated changes in power on the single trial level as explained in the Statistical Analysis section below.

### Drift Diffusion Model (DDM)

To investigate the computational bases of decisions, we fit a multi-attribute DDM (Ratcliff & McKoon, 2008; Smith & Ratcliff, 2004) to the choice and reaction time (RT) data from the SRCT, separately for each subject in each condition. The model included six parameters per condition: three parameters related to value on each trial (weights on tastiness, *w_tastiness_*, healthiness, *w_healthiness_*, as well as a constant *ValConst*), starting point bias (*spbias*), non-decision time (*ndt*) representing post-stimulus perceptual and pre-response motor processes, and decision threshold (*trs*). To identify the best-fitting parameters, we used the differentially-evolving Monte-Carlo Markov Chain method adapted from MATLAB scripts developed by Holmes and Trueblood (2018). In brief, for every trial, we used the DDM to simulate the likelihood distribution of choices and RTs given a specific combination of food attributes and computational parameters, and then used this distribution to construct a Bayesian estimate of the posterior likelihood of the computational parameter values given the data. For each individual and condition, we used 18 chains (i.e. 3 times the number of free parameters), beginning with uninformative priors and constraining parameter values based on previous work and theoretical bounds. All code for fitting the models is available at [OSF website included upon publication]. To estimate the posterior distribution of the each parameter, we sampled 1500 iterations per chain after an initial burn-in period of 500 samples. Best-fitting parameters were computed as the mean for that parameter averaged over the posterior distribution across all chains.

### Model-simulated neural signals

We used the best-fitting subject- and condition-specific parameter values to simulate the expected time course of three signals on each trial: 1 & 2) the expected *attribute value signals* related to tastiness and healthiness; 3) the expected *EA signal* (Figure 2a). Each signal was computed from food onset to response time using individual subject parameters (Figure 2b). For signals related to attributes, the functions on each trial resembled a boxcar with onset immediately following termination of the perceptual part of non-decision time (i.e., the estimated non-decision time parameter minus 80 ms, to account for motor preparation time that occurs before response), offset immediately upon response, and a height equivalent to the attribute value on each trial (Figure 2b; blue and red lines). The EA signal, in contrast, accumulates noisily from immediately after termination of the perceptual non-decision time until the response, with an average slope equal to the integrated value of the choice (Figure 2b; black lines). This approach thus allowed us to construct three simulated time courses: a predicted trial-by-trial time course for the neural representation of the tastiness attribute (tastiness-signal), a predicted trial-by-trial time course for the neural representation of the healthiness attribute (healthiness-signal), and a predicted time course for the accumulated evidence signal (EA-signal) (Figure 2a and b). To produce a more stable prediction, we constructed 1000 simulated datasets for each subject, and took the average across these simulations.

**Figure 2:**
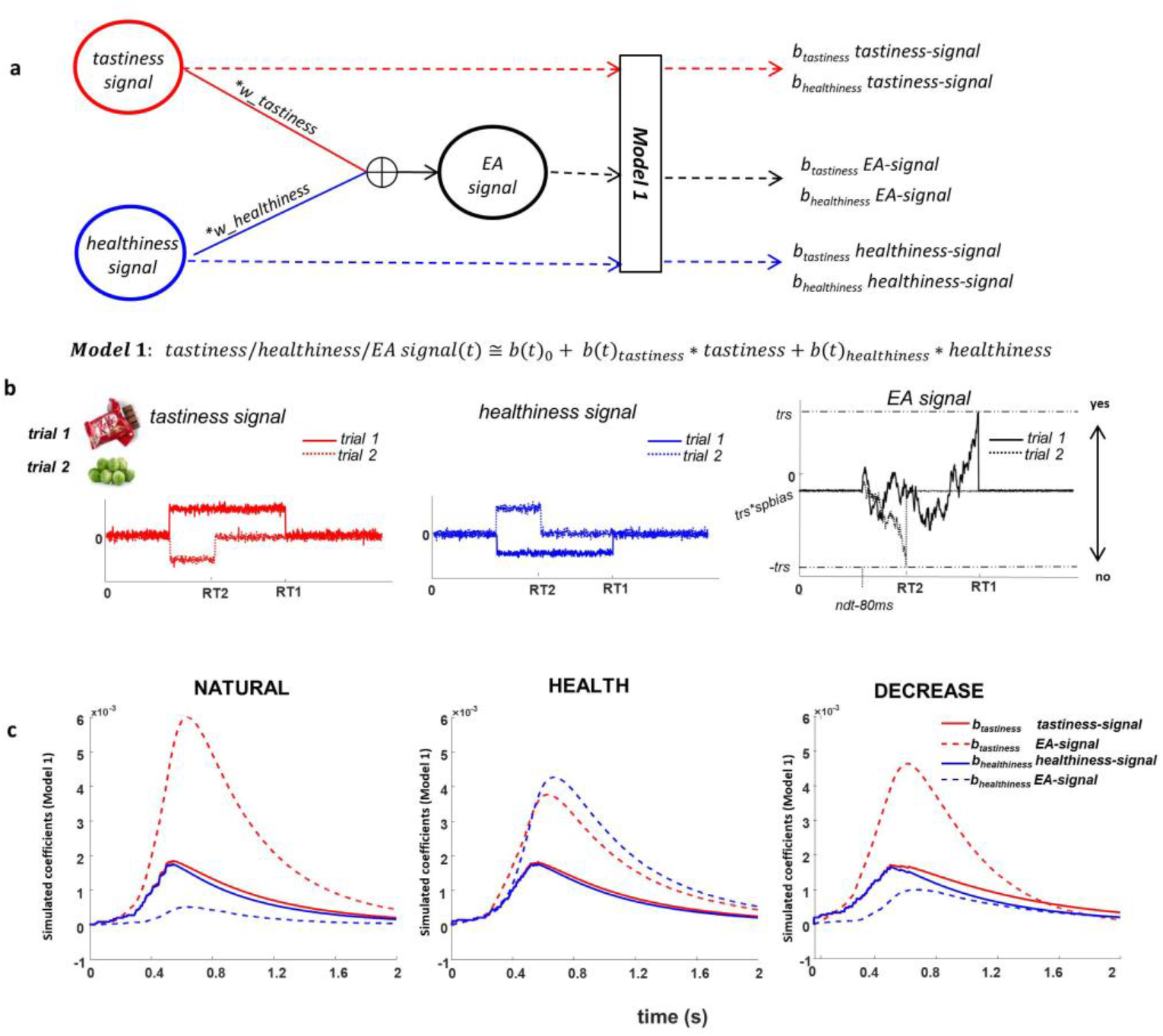
Attribute and evidence accumulation (EA) models. a) Attribute and EA representations based on DDM structure; we assume tastiness and healthiness signals integrate by DDM drift rates and create the EA signal. Model 1 computes the association of individually perceived tastiness and healthiness of each food with each of the tastiness, healthiness, and EA signals. b) examples of time course of tastiness (red), healthiness (blue), and EA (black) signals in 2 trials where a tasty but unhealthy (trial 1, solid lines) and a healthy but non-tasty (trial 2, dotted lines) food were presented. RT1 and RT2 are the reaction times on trial 1 and trial 2. Parameters used for these simulation examples are w_tastiness_ = .07, w_healthiness_ = .01, ValConst = .05, threshold = .09, spbias = −.07, nondec = .462 s c) Model 1 coefficients showing the association between food tastiness ratings and the simulated tastiness signal (solid red line), food tastiness ratings and the simulated EA signal (dashed red line), food healthiness ratings and the simulated healthiness signal (solid blue line) and food healthiness ratings and the simulated EA signal (dashed blue line) using individual DDM parameters in NATURAL, HEALTH, and DECREASE conditions. Coefficients are averaged across subjects.

Finally, we asked how such signals might appear in neural data, when locked to onset of the food. For example, to determine the expected time course of a neural signal representing the tastiness attribute (i.e., tastiness-signal), within each condition (i.e. Natural, Health, Decrease), we computed a separate regression for each time point *t* = 0 to 2000, (i.e. from food onset to 2000 ms afterwards), using the average model-simulated trial-by-trial tastiness signal at that time point as the dependent variable, and the subject’s trial-by-trial tastiness and healthiness ratings as predictors. We repeated this procedure again with the simulated healthiness signal as the dependent variable and with the simulated EA signal as the dependent variable (Model 1).

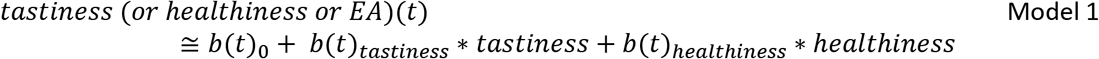

The resulting *b(t)_tastiness_* and *b(t)_healthiness_* coefficients at each time point represent when and to what extent we would expect to observe an association between 1) the tastiness ratings and a neural signal computing tastiness, 2) healthiness ratings and a neural signal computing healthiness, and 3) tastiness and healthiness ratings and a neural signal performing a process of weighted evidence accumulation. The distinct time course of these betas for each condition can be seen in Figure 2c. We used these model-predicted time courses to identify oscillatory EEG signals conforming to model predictions of either attribute or EA signals (see below for details).

### Statistical analysis

#### Behaviour andDDM

We analyzed behavioural data both on the subject and trial level. We used 3-way ANOVAs with NATURAL, HEALTH, and DECREASE as within-subjects factors to study the effect of regulation on RT, choice, and DDM parameters across subjects. Post-hoc paired t-tests were used to compare regulation conditions with NATURAL and with each other.

#### EEG

For comparison to simulated computational time courses (see above), the power of frequencies from 1 to 80 Hz was averaged into six frequency bands: delta (1-3 Hz), theta (4-8 Hz), alpha (9-12 Hz), beta (13-35 Hz), gamma1 (36-59Hz), and gamma2 (61-80 Hz). Power was calculated for each 49ms time bin (25 data points) for all channels. We then regressed trial-by-trial power estimates (*tfPower*) in every frequency band onto the trial-by-trial *tastiness* and *healthiness* of the food item presented on each trial (Model 2).

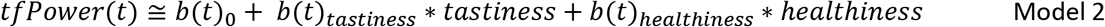

Using this model, we extracted *b(t)_tastiness_* and *b(t)_healthiness_* for each frequency band, time bin, and channel for each subject separately in every condition, allowing us to construct an empirically-observed time course of beta coefficients that was analogous to the model-predicted time courses estimated from Model 1 (see above). We then regressed time courses of *b(t)_tastiness_* and *b(t)_healthiness_* from Model 1 for the tastiness, healthiness and EA signals onto their counterparts in Model 2, across frequency bands and channels. Finally, we calculated the correlation between the time course of these coefficients from Model 1 and Model 2 from 0 – 1s following stimulus presentation (*r_tf-model_*). This allowed us to examine the extent to which the time course of attribute-related components of oscillatory power resembled predictions of the computational model.

We retained only those channel × frequency bands with joint significance criterion where 1) *r_tf-model_* (i.e., the correlation between *b*-coefficients of power from Model 2 and *b*-coefficients of tastiness, healthiness, or EA signals from Model 1 was > .85 and significant at p < .05, Bonferroni corrected for number of comparisons (64 channels × 6 frequency bands) *and* where Model 2 *b*-coefficients of power in the corresponding frequency were significantly different from zero (df = 49, p <.05) across participants within a contiguous window of +/- 100ms around the peak of averaged model predictions for a given signal (i.e., tastiness, healthiness, or EA signal) with a window length cut-off at 1000 ms post-food stimulus.

Additionally, in *post hoc* exploratory analyses, we examined the time course of *b*-coefficients of Model 2 for effects that did not follow model-predicted temporal trajectories. For this analysis, we retained any channels with at least 6 consecutive time bins (i.e. > 300ms) in which *b*-coefficients of power were significantly different from zero (df = 49, p <.05) across participants with a window length cut-off at 1000 ms post-food stimulus.

## Results

### Effects of regulatory focus on behaviour

We first sought to confirm that regulation influenced choice behaviour, focusing first on overall food acceptance rates. A 3-way ANOVA with condition (NATURAL, HEALTH, DECREASE) as a within-subjects factor revealed differences in acceptance rate across conditions (F(2,98) = 18.73, p < .001). As expected, subjects accepted foods less often in DECREASE compared to both NATURAL (*t* (49) = −4.0, *p* < .001) and HEALTH (*t* (49) = −3.9, *p* < .001), which did not differ from each other (*p* > .05). Response times were also affected by regulation (F(2,98) = 7.33, p = .004. Subjects were faster in the NATURAL compared to both HEALTH (*t* (49) = −5.7, *p* < .001) and DECREASE (*t* (49) = −2.6, *p* = .01) which were not significantly different from each other (*p* > .05).

### Effects of regulatory focus on computational model parameters

To examine the computational bases of these effects, we estimated a DDM with six parameters: weights on tastiness and healthiness, a generic value constant (i.e., a general tendency to assign positive or negative values to foods, unrelated to tastiness or healthiness), threshold, non-decision time, and starting point bias (see Methods for details; Figure S1).

As expected, we found that regulation had a significant effect on the weight given to tastiness (*w_tastiness_*: F(2,98) = 26.44, p < .001) and healthiness (*w_healthiness_*: F(2,98) = 51.63, p < .001) and on the value constant (F(2,98) = 10.94, p < .001). Post-hoc paired t-tests showed that weight on tastiness decreased to a similar extent in HEALTH (t (49) = −6.48, p < .001) and DECREASE (t (49) = −5.55, p < 0.001) compared to NATURAL. By contrast, while the weight on healthiness (*w_healthiness_*) significantly increased in both HEALTH and DECREASE compared to NATURAL (Health: t (49) = 8.77, *p* < .001; DECREASE: t(49) = 2.79, p = .007), it was significantly larger in HEALTH compared to DECREASE (t(49) = 6.52, p < .001). In addition, the value constant *(ValConst)* was more negative in DECREASE compared to NATURAL (t (49) = −3.05, p = .004) and HEALTH (t (49) = −4, p < .001), but was not significantly different between NATURAL and HEALTH (p>.05). Thus, while both strategies decreased the influence of tastiness on choice, HEALTH focus specifically increased the influence of healthiness, and DECREASE resulted in a decrease in value that is non-specific to tastiness or healthiness.

Regulation also influenced other parameters of the model beyond attribute weights. In particular, it changed the decision threshold (*trs*: F(2,98) = 6.38, p =.002) and non-decision time (*ndt*: F(2,98) = 9.22, p<.001). Notably, we found little evidence that regulation affected starting point biases (*spbias*: p>.05), suggesting that most of the effects of regulation occurred during post-stimulus processing, rather than in pre-stimulus preparation. Post-hoc paired t-tests on significant effects showed that decision threshold was higher in DECREASE compared to NATURAL (t(49) = 2.83, p =.007) and HEALTH (t(49) = 2.06, p =.04) and in Health compared to NATURAL (t(49) = 5.02, p <.001). Non-decision time was also shorter in DECREASE compared to NATURAL (t(49) = −3.58, p <.001) and HEALTH (t(49) = −3.06, p =.004) but was not different in HEALTH and NATURAL conditions. These results indicate that regulation more strongly changes value components as opposed to creating a value-independent bias.

### Model simulations of expected EA dynamics

Before turning to the neural results, we sought to use the best-fitting model parameters to simulate the expected time course of three neural signals related to evidence and evidence accumulation: the expected time course of a signal specifically encoding tastiness information, the expected time course of a signal specifically encoding healthiness information, and the expected time course of a signal related to the EA process itself, which integrates both tastiness and healthiness (see Methods for details). We then asked how these signals should correlate with the magnitude of tastiness and healthiness ratings across time, essentially replicating the analyses we applied to the observed EEG data.

Importantly, we observed a characteristic profile that distinguished simulated neural signals primarily related to attributes from neural signals primarily related to the EA process. Across the three conditions, the average association between food tastiness ratings and the simulated tastiness-signal (*b_tastiness_* tastiness-signal) reached a maximum at 542 ms, significantly earlier than the corresponding peak of average association between tastiness ratings and the EA-signal (*b_tastiness_* EA-signal) at 628 ms (t(49) = −4.3, p <.001; Figure 2c). Similarly, the average association between food healthiness ratings and the healthiness-signal (*b_healthiness_* healthiness-signal) reached a maximum at 542 ms, significantly earlier than the peak of average contribution of food healthiness to the EA-signal (*b_healthiness_* healthiness-signal) at 660 ms (t(49) = −4.1, p <.001; Figure 2c). These differences in the time course of model simulations suggest that signals related to attribute representations would be expected to peak around 540 ms, while signals related to evidence accumulation would be expected to peak later, around 640 ms.

### Alpha power correlates with the simulated influence of tastiness on evidence accumulation

To compare the model-simulated results against observed neural responses, we extracted trial-by-trial event-related changes in power of all oscillatory ranges from 200 ms before to 1000 ms after the presentation of the food stimulus. We then regressed averaged power in each trial on trial-by-trial tastiness ratings (Model 2), separately for each of 25 time bins beginning 200ms prior to food onset and ending 1000 ms after food presentation (duration ~49 ms). We asked whether there were any oscillatory signatures that correlated with the model-predicted tastiness signal when no self-regulation was applied, and whether self-regulation in HEALTH and DECREASE conditions modulated these signatures.

Based on our significance criteria (see Methods for details), we identified an alpha frequency signature showing a regulation-modulated sensitivity to tastiness that was in line with model predictions. In particular, in the NATURAL condition, we found a significant change in alpha power over left-frontal and right parietal-occipital channels that negatively correlated with tastiness ratings from 536-1000 ms after presentation of food (t(49) = −3.37, p = .004; Figure 3a). By contrast, alpha power during the two regulation conditions (HEALTH and DECREASE) was not significantly associated with tastiness. The difference in correlation between tastiness and alpha power in the window from 536 to 1000ms was significant for the contrasts of both HEALTH vs NATURAL (t (49) = 2.22, p = .03) and DECREASE vs NATURAL (t (49) = 2.35, p = .02; Figure 3c). For simplicity, we call this effect the *“alpha-tastiness correlate* “.

**Figure 3:**
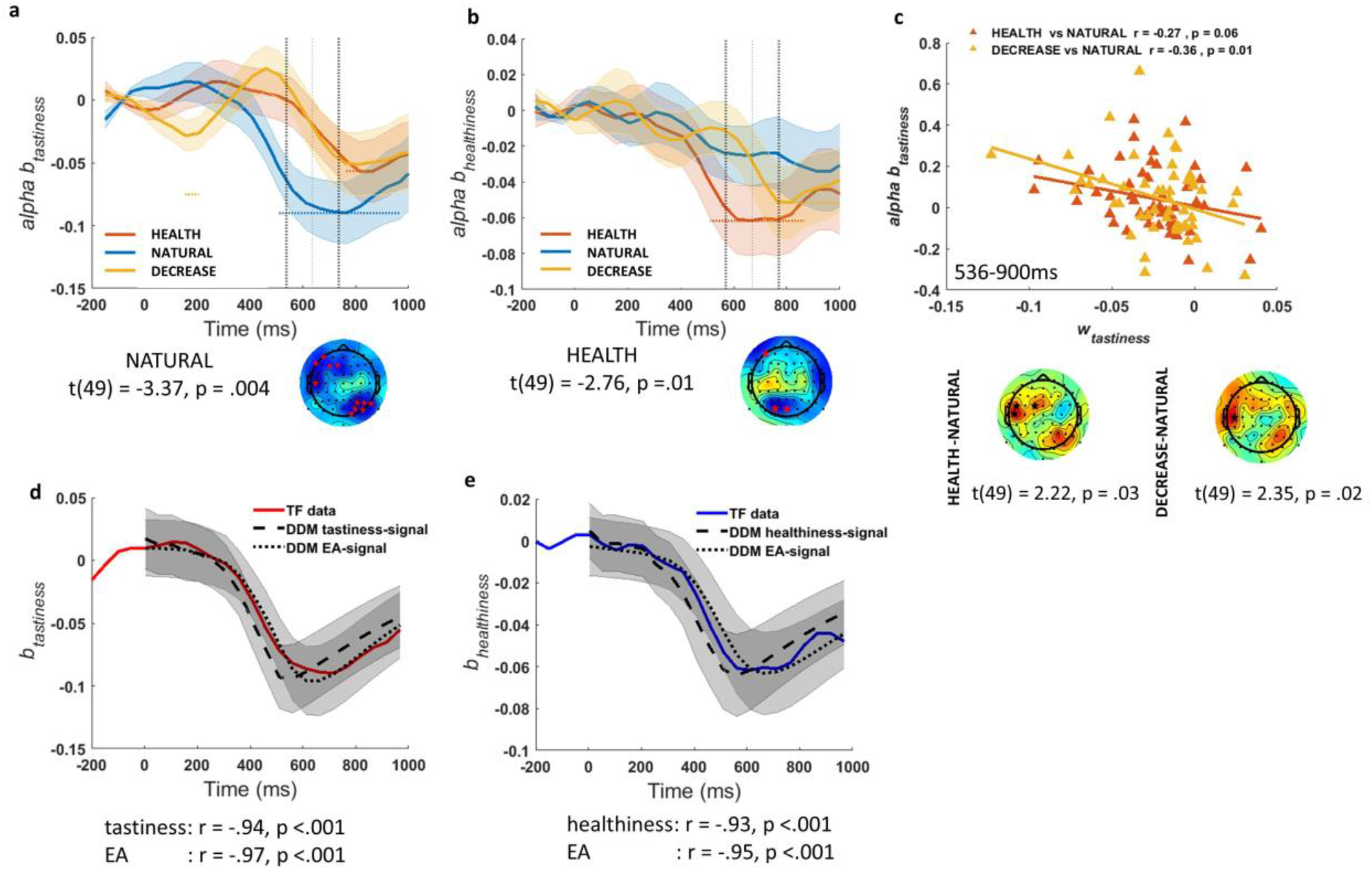
Association of alpha power with tastiness and healthiness. a) alpha-tastiness correlate: alpha power b_tastiness_ in Model 2 in NATURAL condition satisfies significance criteria (see Methods) on frontal and occipital-parietal channels ~530-1000 ms post-food, b) alpha–healthiness correlate: alpha power b_healthiness_ in Model 2 in HEALTH condition satisfies significance criteria (see Methods) on frontal and occipital channels ~550-1000 ms post-food, c) alpha-tastiness correlate predicts successful down-regulation of tastiness influence (*w_tastiness_*) across subjects, d) correlation between the alphatastiness correlate (red line) and tastiness coefficients of Model 1 for tastiness-signal (dashed black line) and EA-signal (dotted black line), e) correlation between the alpha correlate of healthiness (blue line) and healthiness coefficients of Model 1 for healthiness-signal (dashed black line) and EA-signal (dotted black line)

We further asked whether changes in the *alpha-tastiness correlate* in regulation conditions compared to NATURAL predicted individual differences in DDM tastiness weight changes (Δ*w_tastiness_*) in regulation compared to NATURAL, as would be expected if this signal represents a computation directly related to choice. As hypothesized, we found a correlation between the decrease in the weight on tastiness (Δ*w_tastiness_*) and the decrease in the *alpha-tastiness correlate* ~530-1000 ms post-food (starting 100 ms prior to model peak latency), for both the comparison of DECREASE vs NATURAL (r = −. 36, p = .01) and HEALTH vs NATURAL (r = −.27, p = .06), although the latter correlation fell just short of significance (Figure 3c).

Finally, we tested whether alpha power in channels where alpha-tastiness correlates were found correlated better with the attribute or EA signals predicted by the DDM (Figure 2c). Importantly, we found that the *alpha-tastiness correlate* was predicted closely by both the tastiness signal and the EA signal (Figure 3d), with the EA signal showing a more precise relationship across time. However, given the high correlations between both signals, we refrain from making strong inferences about the precise source of this signal. Regardless, these results strongly suggest that regulation operates in part by altering alpha frequency representations of tastiness attributes.

### Alpha power correlates with the simulated influence of healthiness on evidence accumulation

Next, we turned to the healthiness attribute, using a similar analytical approach. Notably, in the HEALTH condition, we again found significant effects restricted to alpha power in frontal and parietal-occipital channels. Alpha power in these channels was negatively correlated with healthiness from ~550-1000 ms after presentation of food (t(49) = −2.76, p = .01; *alpha-healthiness correlate*; Figure 3b). Similar to observations regarding tastiness, the timing of this signal correlated with the predicted time course of model signals, and did so more strongly for the EA signal than the healthiness signal (Figure 3e).

As might be expected from model predictions, alpha power during the NATURAL and DECREASE conditions was not significantly associated with healthiness, although the contrast of average *b_healthiness_* for alpha power between HEALTH and DISTNCE or NATURAL in these channels ~550-1000 ms post-food failed to reach significance. Unlike the *alpha-tastiness correlate*, we found no evidence that the differences in *alpha-healthiness correlate* across conditions correlated with individual differences in the change in model parameters for the healthiness weight.

### Early theta power correlates with later suppression of tastiness influence on EA

Having shown that we observed a striking correspondence between model predicted changes in attribute representations and suppression of oscillations in the alpha range, we next turned to a more exploratory analysis. Specifically, we asked whether there were any oscillatory changes that might distinguish regulatory conditions in relation to *tastiness* and *healthiness*, but that did *not* conform to model predictions. Strikingly, and in contrast to alpha power, which tracks tastiness in the NATURAL condition but not regulation conditions, we found a single pattern matching our significance criterion: theta power at channels over parietal-occipital and frontal areas was positively correlated with tastiness early in the trial (150-500ms) in both HEALTH (t(49) = 3.28, p = .005; Figure 4a) and DECREASE (t(49) = 3.48, p = .004; Figure 4b) conditions independently. This effect was also larger compared to NATURAL in both HEALTH (t(49)= 3.01, p = .01; occipital channels) and DECREASE (t(49) = 2.29, p = .03; occipital and left central channels) (Figure 4c). For simplicity, we call this effect “*theta-tastiness correlate*”.

**Figure 4:**
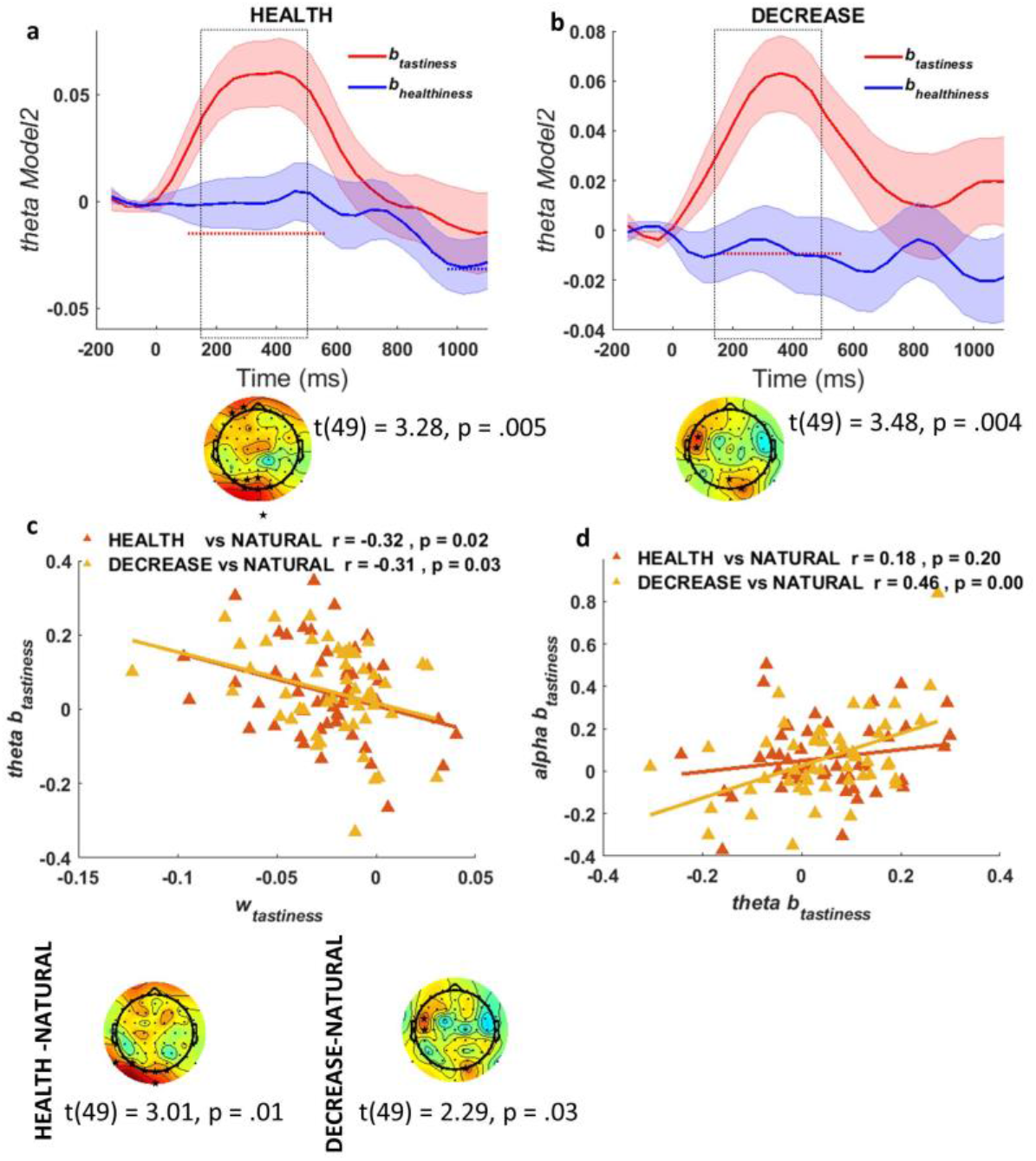
Theta-tastiness correlate: theta power coefficients for tastiness in Model 2 satisfy significance criteria (see Methods) at frontal and occipital channels in a) HEALTH and b) DECREASE conditions ~150-500 ms post-food. Theta correlate of tastiness predicts c) successful down-regulation of tastiness influence (*w_tastiness_*) and is correlated with d) alpha-tastiness correlate across subjects

Given the early appearance of this signal compared to the *alpha-correlate of tastiness*, we speculated that it might represent a “control” process, rather than a representation of the tastiness attribute per se. This interpretation predicts that, despite correlating with tastiness, the magnitude of this correlation should actually be *negatively* associated with regulation-induced decreases in the weight on tastiness, and that theta and alpha signatures might also be related. These predictions were strongly confirmed. We found that subjects with greater increases in the *theta-tastiness correlate* between 150-500ms showed greater *decreases* in the weight on tastiness (Δ*w*_tastiness_), and this was independently true for the contrast of both HEALTH (r = −.34, p = .02) and DECREASE (r = −.31, p = .03) vs NATURAL (Figure 3c). Moreover, changes in the *alpha-tastiness correlate* and the *theta-tastiness correlate* in DECREASE vs NATURAL were correlated across subjects (r = .46, p =.001; Figure 3d). In other words, the more strongly tastiness was encoded in theta power early in the trial, the *less* strongly it was represented in the suppression of alpha power later in the trial. A similar though non-significant relationship was observed in the HEALTH condition (r = .18, p =.2; Figure 3d).

No other frequency ranges showed any effects of attribute or condition that met our criteria.

## Discussion

How does cognitive self-regulation alter the dynamic process of value computation and evidence accumulation? Do different strategies operate via different mechanisms? Using a computational model of attribute valuation and evidence accumulation, we found that alpha power recorded from frontal and occipital areas of the scalp tracks variation in the tastiness of food when subjects naturally decide what foods to eat: the tastier the food, the lower the alpha power. When subjects regulate their responses in a way that reduces the behavioural influence of tastiness, this function is compromised, i.e., alpha no longer tracks tastiness values. In addition, when subjects focus on healthiness, alpha in similar regions tracks variation in healthiness of food: the healthier the food, the lower the alpha power. The time course of both the tastiness and healthiness effects in alpha power followed closely with the predicted time course of EA in our computational model, and predicted individual differences in successful downregulation of the influence of tastiness on choice across two different regulatory strategies. In addition, we found an earlier rise in theta power at frontal and occipital areas that, while it did not correlate with model-predicted time courses, did correlate with food tastiness when subjects regulated their decisions: the tastier the food, the higher theta power. In contrast to alpha power correlates however, correlation with tastiness in early theta power predicted *decreases* in the influence of food tastiness on subjects’ choices. This was true for both regulatory strategies. These findings suggest that regulation may recruit early theta-mediated control processes to modulate the influence of attributes on alpha suppression during evidence accumulation.

### Functional significance of alpha suppression in EA

Suppression of alpha oscillations, also called the event-related desynchronization of alpha, is widely known to reflect active information processing involving perception and attention and appears to be driven by cortico-cortical networks across frontal and parietal brain regions (Foxe & Snyder, 2011; Klimesch, 2012; Klimesch, Sauseng, & Hanslmayr, 2007; Samaha, Boutonnet, Postle, & Lupyan, 2018). The degree of alpha suppression also predicts success in tasks where a higher level of cortical excitation is required (Klimesch et al., 2007), while inducing transient changes in alpha activity by external stimulation leads to predictable changes in cognitive performance (Klimesch, Sauseng, & Gerloff, 2003). This literature strongly supports our observation that alpha suppression correlated with food attributes, i.e., with tastiness when deciding naturally and with healthiness when focusing on health, and also correlates with behavioural evidence of regulatory success (Figure 3 a, b, c).

Other work also supports the idea that prefrontal and posterior alpha suppression correlate with EA, although such studies have focused on perceptual decision making (Kloosterman et al., 2019; vanVugt et al., 2012; Werkle-Bergner et al., 2014). This work suggests that alpha suppression may relate to the control of cortical excitability in relation to accumulation of sensory evidence (Kloosterman et al., 2019). Yet the precise role of alpha suppression in representation of *value* during EA is not clear. Here, we found a striking similarity between the time course of alpha correlates of tastiness and healthiness and the representation of these attributes in the simulated neural signals predicted by the best-fitting parameters of the DDM, particularly the EA signal (Figure 3 d, e).

In this respect, it is interesting to speculate on the meaning of differences we observed between the relationship of alpha suppression to the influences of tastiness and healthiness on behaviour. While the *alpha-tastiness correlate* showed a strong presence in occipital and frontal areas and correlated independently in both regulatory conditions with individual differences in down-regulation of tastiness, the *alpha-healthiness correlate* was comparatively less consistent: It showed a more limited presence in posterior and frontal regions and did not correlate with up-regulation of healthiness across subjects. It is possible that this is simply due to lack of power to detect significant findings. Alternatively, we speculate that the HEALTH condition may activate both down-regulation of tastiness and up-regulation of healthiness. In combination with the relative *dominance* of tastiness attribute, this joint set of effects could result in a weaker alpha-healthiness correlate. Overall, the close relationship between the DDM time course and alpha suppression indicates a quantifiable role for alpha suppression and its disruption during self-regulation.

Our results also indicated that alpha correlates of tastiness and healthiness corresponded more closely with the predicted time course of an EA signal as opposed to the DDM attribute signals themselves. Interestingly, we observed no specific signature that fully matched the predictions of our attribute correlations. Although speculative, this might indicate that EEG alpha oscillations, representing an EA process, might be more consistent across subjects, while attribute processing might be more heterogeneous. Indeed, both healthiness and tastiness might themselves be “composite” attributes constructed from a number of component attributes (e.g., sweetness, saltiness, texture might all contribute to tastiness; calories, nutrients, and/or fiber might all contribute to subjectively perceived healthiness). Alternatively, attribute representations might show variable temporal dynamics across trials and subjects, defying assumptions in our model about their time course (see Figure 2b). Future work will be needed to determine whether EEG might be used to inform a more nuanced, time-dependent DDM (Maier, Raja Beharelle, Polanía, Ruff, & Hare, 2020).

### Functional significance of theta in self-regulation

Over the past several years, a consensus has emerged in the field that theta oscillations in the frontal cortex reflect cognitive control and integration of resources for cognitive control (e.g., Cavanagh & Frank, 2014; Mas-Herrero & Marco-Pallarés, 2016). Midfrontal theta, in particular, activates in response to any informative event or stimulus that conveys a need for behavioural adjustment or shift of attention (Cavanagh & Frank, 2014) and recruits regions underlying different cognitive operations to realize this need for control (Mas-Herrero & Marco-Pallarés, 2016). Posterior theta oscillations, on the other hand, may instead reflect the encoding of new information by the hippocampus in several memory operations (Axmacher et al., 2010; Klimesch, 1999; Scholz, Schneider, & Rose, 2017; Staudigl & Hanslmayr, 2013; Tesche & Karhu, 2000).

By contrast, evidence for the functional significance of the lateral frontal and posterior theta oscillations we observed in this task has not converged on a unified account. There is scattered evidence that early parietal and occipital theta oscillations correlate with value in high conflict choices (Hunt et al., 2012) and with cognitive load during mental calculation and multi-tasking (Wang, Jung, & Lin, 2018). On the other hand, frontal theta in value-based decisions also shows sensitivity to emotional stimuli and efficient sensory integration in different studies (Cavanagh et al., 2011; Nayak, Kuo, & Tsai, 2019; Tosun, Berkay, Sack, Çakmak, & Balci, 2017; Zavala et al., 2016).

Our findings may help to shed light on these inconsistencies. Notably, the *theta-tastiness correlate* in our study appeared in both regulation conditions (but not unregulated decisions) around the same time, with a shared occipital component, and a similar though not identical frontal component (although the rough spatial resolution of EEG prevents us from making strong conclusions about whether such differences imply distinct neural sources). This signal was sensitive to tastiness in both regulation conditions, but not to healthiness while focusing on health. In other words, we observed stronger theta signals on precisely those trials which might be hardest to regulate (i.e., trials with tastier foods). This suggests that the *theta-tastiness correlate* may represent more of an inhibitory control mechanism as opposed to an attentional shift to a new attribute. The fact that this signal was observed independently in both regulatory conditions suggests that it represents a common operation in self-control, independent of the specific strategy/goal.

What might the mechanisms of such theta-mediated attribute suppression be? Based on the proposed role for theta in signaling the need for control and implementation of control (Cavanagh & Frank, 2014), we speculate that, when subjects encounter tastier foods in trials where they need to apply self-control, theta oscillations communicate a need for changes in sensory processing of the stimulus. As the need for control diminishes in the case of less tasty foods, or when self-control is not required, theta may also decrease. Given the posterior component of theta, an important question for future studies will be to investigate whether it influences perceptual processing or memories of food items that are presented in the regulatory conditions as a function of their tastiness (Shadlen & Shohamy, 2016). Future studies using brain stimulation should also investigate whether disruption of early theta power causes impairments in regulation. Another interesting question for further studies will be to address whether theta correlates of tastiness can be useful in predicting the dynamics of evidence accumulation by sequential sampling models such as DDM.

### Comparisons to previous work on value and regulation

Our findings contribute to a growing but still scant EEG literature on the neural representation of components of value in the EA process. While some studies have examined modulation of EEG during self-control (Harris et al., 2013), and other studies have examined oscillatory correlates of EA itself (Hunt et al., 2012; vanVugt et al., 2012; Werkle-Bergner et al., 2014), ours is the first study to date to combine the two approaches. In this respect, it is informative to compare our results to previous work. For example, Harris et al. (2013) reported an early attentional ERP component in occipital areas (N170) that was enhanced in successful self-control and later centro-parietal ERP components localized to the ventromedial prefrontal cortex that were sensitive to the overall value and tastiness and healthiness attributes. Similarly, in our study, we found two temporally distinct oscillatory signatures corresponding to valuation (alpha suppression around 600ms post-stimulus) and regulation (theta enhancement around 200ms post-stimulus). While a direct comparison of these two sets of findings is not possible due to differences in the study design and analysis, we speculate that the early EEG predictors of regulation observed in our study as well as in Harris et al. (2013) might indicate a common oscillatory mechanism (occipital theta power), that controls the processing of attributes in the service of regulation. We also speculate that the later components of attribute value represented by alpha suppression in our study reflects the *controlled* integration of sensory evidence by areas involved in decision making and response selection.

Importantly, while we do observe neural signatures consistent with EA signals in our study, our results fail to replicate some previous reports in the literature finding a distinct gamma (46-64Hz) signature of value-based EA (Polanía et al., 2014) at parietal and fronto-polar sites prior to response. Intriguingly, gamma phase coherence in this prior study revealed fronto-parietal connectivity during value-based decisions and not perceptual decisions. To test specifically whether we replicated this finding, we examined oscillatory power in the NATURAL condition averaged over all channels and specifically at Fz and Pz channels, but did not find an enhancement of gamma in relation to our pre-food 200 ms baseline (Figure S2a). To test whether this is due to higher variability in gamma phase in food-locked analysis, we also tested response-locked oscillatory power in relation to the pre-food baseline, but failed to find a significant increase in gamma power (Figure S2b).

We speculate that these discrepancies may be due to differences in the cognitive and memory demands presented by the different task paradigms. For example, our subjects make an accept/reject response to one food stimulus whereas in Polania et al. (2014), they responded by choosing one of two food stimuli presented on the upper or lower side of the screen while a scrambled version of each image was displayed by their side (Figure 1A in Polania et al., 2014). This more complex paradigm might elicit higher frequency responses due to differences in visual and spatial processes. Alternatively, it is possible that the choice between two items vs accepting or rejecting a single item, although behaviourally comparable in terms of the decision value, might require distinct working memory processes. Future work directly comparing different modes of choice (e.g. single-item vs. binary or multi-item choice) will be needed to resolve some of these questions.

In addition to gamma signals, Polania et al. (2014) found that fronto-central low-beta (18-20 Hz) power was also negatively correlated with EA. Similarly, we also observed suppression of low-beta power (Figure S2), though to a lesser degree compared to alpha power. A full match to the single-trial value-based predictions of our DDM model regarding EA only occurred with alpha suppression (Figure 3 d,e), and did not match responses in the beta range. Notably, Polania et al. (2014) did not examine frequencies below 15Hz, or look for the kind of trial-by-trial variation in value that we identify here, making a direct comparison of our results to theirs difficult.

### Conclusion

Taken together, our results highlight the potential for model-based analyses to lend new insight into the neural mechanisms underlying self-regulation in value-based choice. Our findings corroborate previous accounts suggesting that alpha suppression tracks EA, and further suggest that alpha suppression may represent an identifiable signature indicating the influence of value-based attributes during both regulated and unregulated choice. Our study is also the first to our knowledge to find evidence that theta oscillations might play a role in inhibiting the influence of attribute values on evidence accumulation, independent of regulation strategy. These findings point to the importance of understanding the distinct neural mechanism by which this inhibition occurs, and may represent a target of future work to enhance the effectiveness of self-regulation in dietary choice.

## Acknowledgements

This study was supported by Natural Sciences and Engineering Research Council of Canada Discovery Grant and the University of Toronto Connaught New Researcher Award.

---

**Figure S1:**
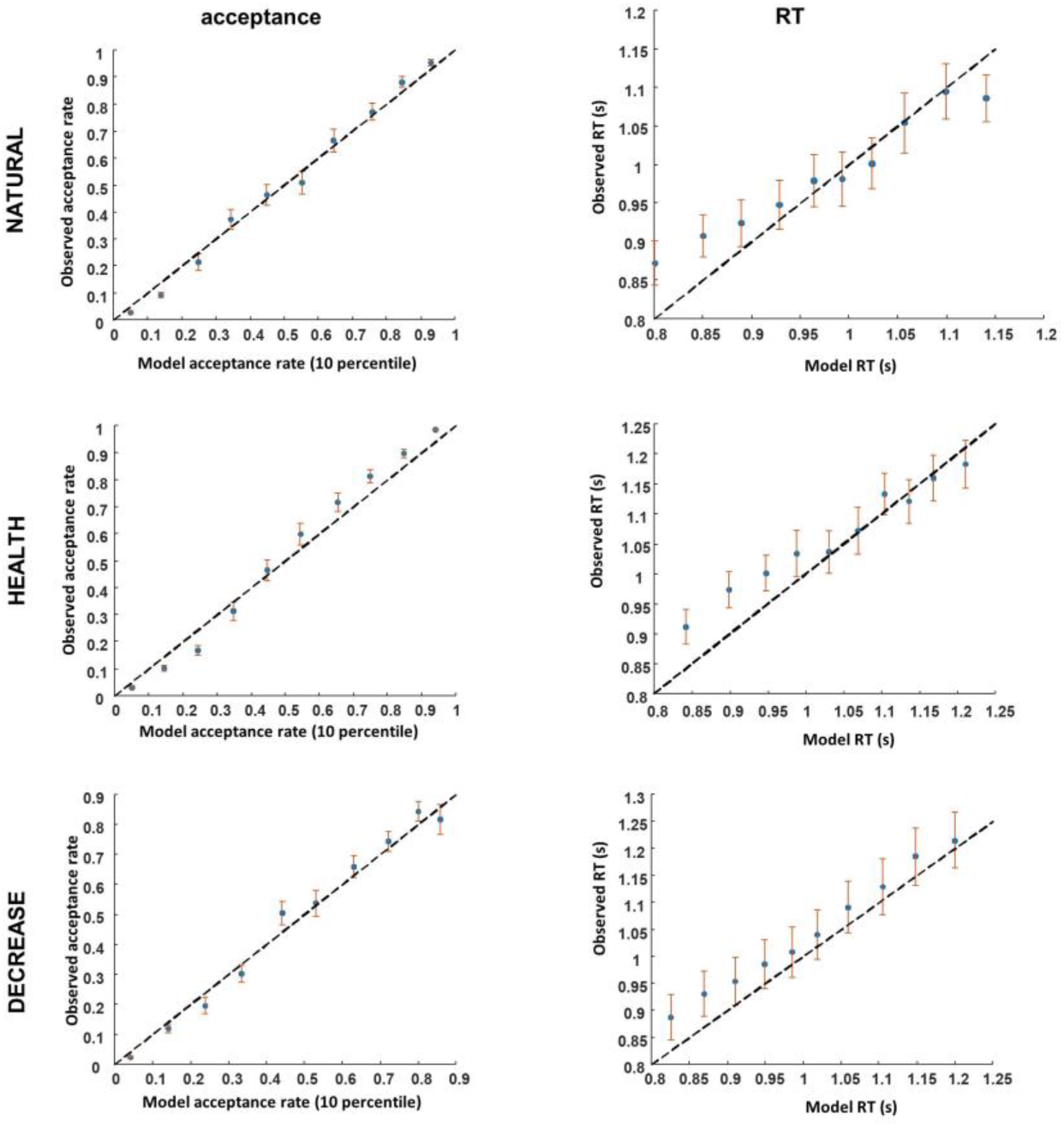
Averaged acceptance rate and RTs simulated by DDM fitted parameters correlate with acceptance rate and RT data from subjects

**Figure S2:**
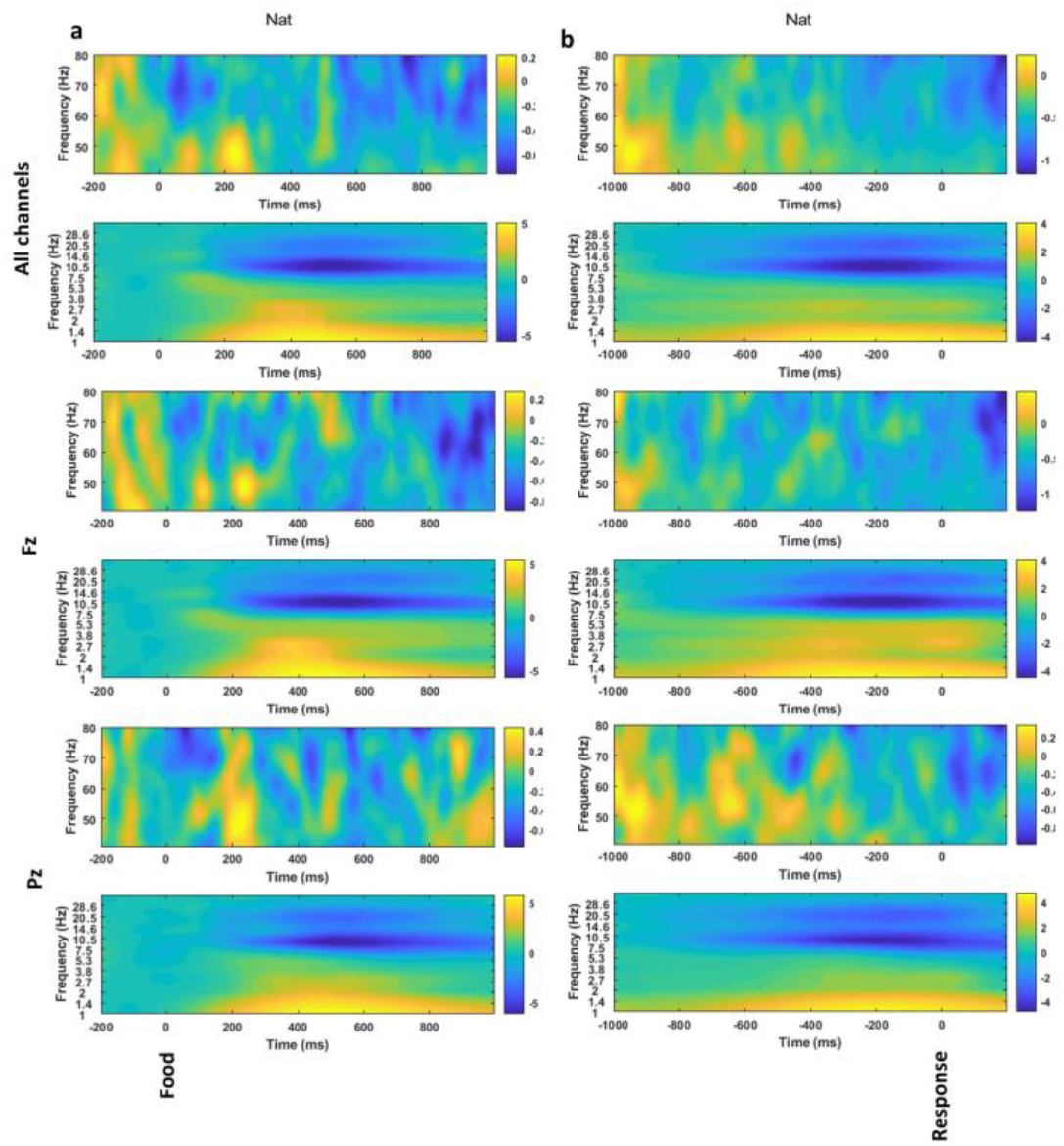
Time frequency plots of averaged channels (top), Fz (middle) and Pz (bottom) in (a) food-, and (b) response-locked EEG

